# The *Pseudomonas aeruginosa* T3SS can contribute to traversal of an *in situ* epithelial multilayer independently of the T3SS needle

**DOI:** 10.1101/2025.01.28.635263

**Authors:** Eric Jedel, Daniel Schator, Naren G. Kumar, Aaron B. Sullivan, Arne Rietsch, David J. Evans, Suzanne M. J. Fleiszig

## Abstract

Multilayered epithelia lining our tissue surfaces normally resist traversal by opportunistic bacteria. Previously, we developed a strategy to experimentally perturbate this resistance *in situ* in the corneas of mouse eyes and used it to show that traversal of a multilayered epithelium by *Pseudomonas aeruginosa* requires ExsA, the transcriptional activator of its type 3 secretion system (T3SS). Here, we developed a novel strategy for quantitively localizing individual traversing bacteria within the *in situ* multilayered corneal epithelium and explored contributions of T3SS components. The results showed that T3SS translocon and T3SS effector mutants had reduced epithelial traversal efficiency. Surprisingly, a Δ*pscC* mutant unable to assemble the T3SS needle traversed as efficiently as wild-type *P. aeruginosa*, while a Δ*exsD* mutant ‘constitutively on’ for T3SS expression was traversal defective. Dispensability of the T3SS needle for effector-mediated traversal was confirmed using a mutant lacking the T3SS operon except the effector genes (Δ*pscU-L* mutant). That mutant reacquired the ability to traverse if complemented with rhamnose-inducible *exsA*, but not if the effector genes were also deleted (Δ*pscU-L*Δ*exoSTY*). Western immunoblot confirmed ExoS in culture supernatants of rhamnose-induced *exsA*-complemented Δ*pscU-L* mutants lacking all T3SS needle protein genes. Together, these results show that epithelial traversal by *P. aeruginosa* can involve T3SS effectors and translocon proteins independently of the T3SS needle previously thought essential for T3SS function. This advances our understanding of *P. aeruginosa* pathogenesis and has relevance to development of therapeutics targeting the T3SS system.

**Importance:** While the capacity to cross an epithelial barrier can be a critical step in bacterial pathogenesis, our understanding of mechanisms involved is derived largely from cell culture experimentation. The latter is due to practical limitations of *in vivo*/*in situ* models and challenge of visualizing individual bacteria in the context of host tissue. Here, factors used by *P. aeruginosa* to traverse an epithelial multilayer *in situ* were studied by: 1) leveraging the transparent properties and superficial location of the cornea, 2) using our established method for enabling bacterial traversal susceptibility, and 3) developing a novel strategy for accurate and quantitative localization of individual traversing bacteria *in situ*. Outcomes showed that T3SS translocon and T3SS effector proteins synergistically contribute to epithelial traversal efficiency independently of the T3SS needle. These findings challenge the assumption that the T3SS needle is essential for T3SS effectors or translocon proteins to contribute to bacterial pathogenesis.

## Introduction

*Pseudomonas aeruginosa* is a common cause of opportunistic infections, able to cause disease in many body sites including the airways, GI tract, skin, urinary tract, eye, brain, heart and blood (1, 2). Susceptibility can be associated with wounds, burns, immunocompromise, chemotherapy, surgery, predisposing diseases (e.g. cystic fibrosis), and use of indwelling medical devices (3–5). In the eye, *P. aeruginosa* is a leading cause of corneal infection (microbial keratitis), estimated to cause vision loss in approximately 2 million people worldwide each year (6–8). A major risk factor for keratitis is contact lens wear (8–10) and *P. aeruginosa* is the most frequently isolated pathogen from lens-associated infections (6, 10). Recently, an outbreak of multidrug resistant *P. aeruginosa* infections was reportedly transmitted *via* simple artificial tears eyedrops, causing infections of the eye and beyond. These infections resulted in multiple instances of vision loss and several deaths, in some instances without contact lens wear or prior eye disease (11–13).

Type three secretion systems (T3SSs) are among various tools Gram-negative bacteria use to export factors across their otherwise impermeable cell membranes. A T3SS is additionally able to inject effector toxins across host cell membranes into the cytoplasm of a target host cell to alter its biology. *P. aeruginosa* encodes a single T3SS that has been shown to make significant contributions to virulence during acute infections of the cornea (14–17) and other body sites (18–22). The *P. aeruginosa* T3SS is encoded by 42 known genes encoding proteins of a multimeric needle core, a translocon pore, and one or more effector toxins (23). ExsA is the only known transcription factor for these genes. Effectors of the *P. aeruginosa* T3SS, like those of other Gram-negative bacteria, can have a multitude of effects on host cells, including modifying their biology to support an intracellular lifestyle (24–27).

All environment-exposed body surfaces are covered by multilayers of epithelial cells, that when healthy resist traversal by *P. aeruginosa* and other opportunistic bacteria. This includes the cornea of the eye, supremely capable of preventing bacterial colonization through a repertoire of intrinsic defenses, some inherent within the corneal epithelium, others conferred by other factors present at the ocular surface (28–34). The cornea’s unique efficacy in this regard makes the corneal surface our only environmentally exposed body surface devoid of a viable bacterial microbiome (35, 36).

If *P. aeruginosa* does adhere to the cornea, which can occur after superficial injury, additional barriers prevent it from traversing the multilayered epithelium to reach the underlying stroma, access to which is required for the initiation of infectious pathology (i.e. keratitis) (28, 37–41). Thus, infection susceptibility requires alterations to the epithelial barrier beyond superficial injury. While that can be accomplished by full thickness injury (42, 43), *P. aeruginosa* corneal infection is most commonly associated with contact lens wear, which predisposes the epithelium to bacterial traversal more subtly by mechanisms not yet well understood (44, 45).

A plethora of studies have been done to explore how *P. aeruginosa* crosses a cultured epithelial cell layer *in vitro*, using a variety of epithelial cell types. For example, airway epithelial cells or MDCK cells were used to show roles for elastase, exotoxin A, type 4 pili, flagella, and the T3SS (46–50). Our own *in vitro* studies using cultured corneal epithelial cells revealed roles for proteases (41), type 4 pilus-associated twitching motility (51), and the T3SS effector ExoU; the latter encoded by only a subset of *P. aeruginosa* strains (cytotoxic strains) and which enables traversal by killing epithelial cells (52). Recently, a study using *in vitro* grown organoids of human airway epithelial cells showed that both the T3SS and type VI secretion system (T6SS) can contribute to *P. aeruginosa* “translocation” (traversal) of an epithelial barrier *via* goblet cell invasion (53).

Thus, our current understanding of how *P. aeruginosa* (and other bacteria) traverse susceptible epithelial layers is based largely on *in vitro* cell culture study outcomes wherein bacteria and host cells are studied in isolation from other factors normally found in their environment *in vivo*. Yet our prior studies have shown that *in vivo* factors modify how bacteria interact with cells, including basement membranes, mucosal fluids, other cell types, nerves, soluble factors, and environmental conditions specific to the tissue site (28, 32, 37, 39, 41, 54, 55). Other limitations of the *in vitro* literature on this topic are that most studies used epithelial cell monolayers that differ from *in vivo* multilayers that contain layers with cells in multiple states of differentiation. Further, transformed cells used by many studies are generally locked into one state of differentiation irrespective of whether they can polarize correctly *in vitro* (e.g. MDCK cells, HeLa cells).

Factors that have hindered development of *in vivo*/*in situ* models for studying epithelial traversal by *P. aeruginosa* and other bacteria include deliberately bypassing epithelial and other tissue barriers to enable infection (e.g. by wounding or bacterial injection through them). Moreover, introducing bacteria *in vivo/in situ* can trigger inflammation that can break down subsequent barriers to bacterial dissemination, reducing or even eliminating the need for bacteria to contribute. Another obstacle to an *in vivo/in situ* study if individual bacteria need to be localized is the challenge of visualizing them in the context of host tissue.

Our published studies have shown that pretreatment of *in vitro* grown corneal epithelial cells with mucosal fluid (tear fluid) increased their resistance to *P. aeruginosa* adhesion, invasion, and cytotoxicity, and increased their traversal resistance when grown as multilayers (39, 56, 57). This was accompanied by profound changes to the epithelial cell’s transcriptome (57). Tear fluid exposure also changes gene expression in *P. aeruginosa*, including multiple genes involved in virulence and survival, reducing its ability to traverse cultured corneal epithelial cells – without directly impacting bacterial viability (39, 57, 58). Other *in vivo* factors that are not present in epithelial cell cultures can modify epithelial-microbe interactions. In the corneal epithelium they include nerves and immune cells, both of which contribute to maintaining epithelial cell homeostasis while also directly recognizing and responding to microbes (28, 31, 32, 59–61). Thus, while cell culture studies have provided a good starting point for understanding epithelial barrier function and mechanisms by which *P. aeruginosa* traverses epithelial cells, *in vivo/in situ* studies are warranted.

Some of our published studies have attempted to address this knowledge gap. In one study, we enabled traversal susceptibility *in vivo* in mice by scratching through the corneal epithelium, then waiting for the time point at which the cell multilayer was reestablished but still remained permissive to bacterial traversal, which occurred 6 hours after scratching (38). Bacterial location and inoculation was quantified in fixed and stained tissue sections. The results showed that mutants lacking ExsA, the transcriptional activator for the T3SS, were unable to traverse the epithelium (38). Since wound healing might have played into the outcome, we later performed a second study enabling susceptibility using more gentle superficial injury, by gently blotting the surface with a KimWipe™, then EGTA-treatment prior to inoculation (37). Using this strategy, we found that bacteria traversing the epithelium did not penetrate past the basal lamina, which when intact functions as a non-specific size exclusion filter (37, 41). This avoids bacterial entry into the stroma and therefore subsequent pathology/inflammation that could complicate studies of how bacteria overcoming epithelial barrier function. In a later study, the blot/EGTA model was used to quantify *P. aeruginosa* traversal of murine corneal epithelium by combining 3D confocal imaging to identify fluorescent bacteria with confocal reflectance microscopy to image the non-fluorescent cornea (62). Results again showed that ExsA was required for *P. aeruginosa* to traverse the corneal epithelium when the mouse was immunocompetent (62).

A limitation of both of our prior studies was that neither method accounted for the complex shape of the cornea, which is curved rather than flat. The problem this presents is that at any plane of imaging the back and front surfaces of the corneal epithelium are not in line. This made it difficult to accurately determine depth of penetration across the population and ascertain whether T3SS mutants were penetration defective, or simply less efficient at traversing the epithelium.

Here, we report development of an advanced analytical imaging approach allowing for precise metrics of location to be determined for all individual bacteria within the curved multilayered epithelium *in situ*. Use of these methods showed that the T3SS impacts penetration/traversal efficiency of *P. aeruginosa* rather than being absolutely required, with roles played by both the T3SS translocon and T3SS effector proteins. Unexpectedly, results showed that the T3SS needle was dispensable for this phenotype, and actually interfered when constitutively expressed.

## Results

### Transcription of the T3SS promotes *P. aeruginosa* traversal efficiency in the multilayered corneal epithelium *in situ*

Our prior study showed that the transcriptional activator of the T3SS (ExsA) contributed to *P. aeruginosa* traversal of the mouse eye’s corneal epithelium (62). Here, we leveraged recent advances in imaging and image analysis to obtain more detail about how ExsA impacts bacterial location, taking into account the complexities of the cornea’s shape while more accurately assessing location of individual members of the traversing population.

As in our previous study, we used confocal reflectance microscopy (CRM) to image the corneal epithelium and GFP-expression to detect bacteria. Raw confocal images were then imported into Imaris software v9.9 for processing, the CRM signal pseudo-colored red (Fig. 1A). As detailed in the methods section, the reflectance signal from the stroma was manually excluded, and a “Surface” representing only signal from the epithelium was generated (Fig. 1B). Apical and basal boundaries of the epithelium were then identified (Fig. 1C). Individual bacteria were identified as a “Spot”, creating objects in the image that each represented one population member (Fig. 1D). The distance (in microns) from the apical and basal boundary of the epithelium was measured for each bacterial “Spot”, and the depth of penetration of each bacterium was calculated as a percentage to provide its relative position in between the upper and lower epithelial boundaries, with 0% being surface adherent and 100% meaning that the bacterium reached the underlying basal lamina (Fig. 1E). Another metric quantified was the proportion of the population able to penetrate beyond the 50% (midpoint) normalized to the thickness of the epithelium in that region of the tissue (as in Fig. 2B).

**Figure 1.**
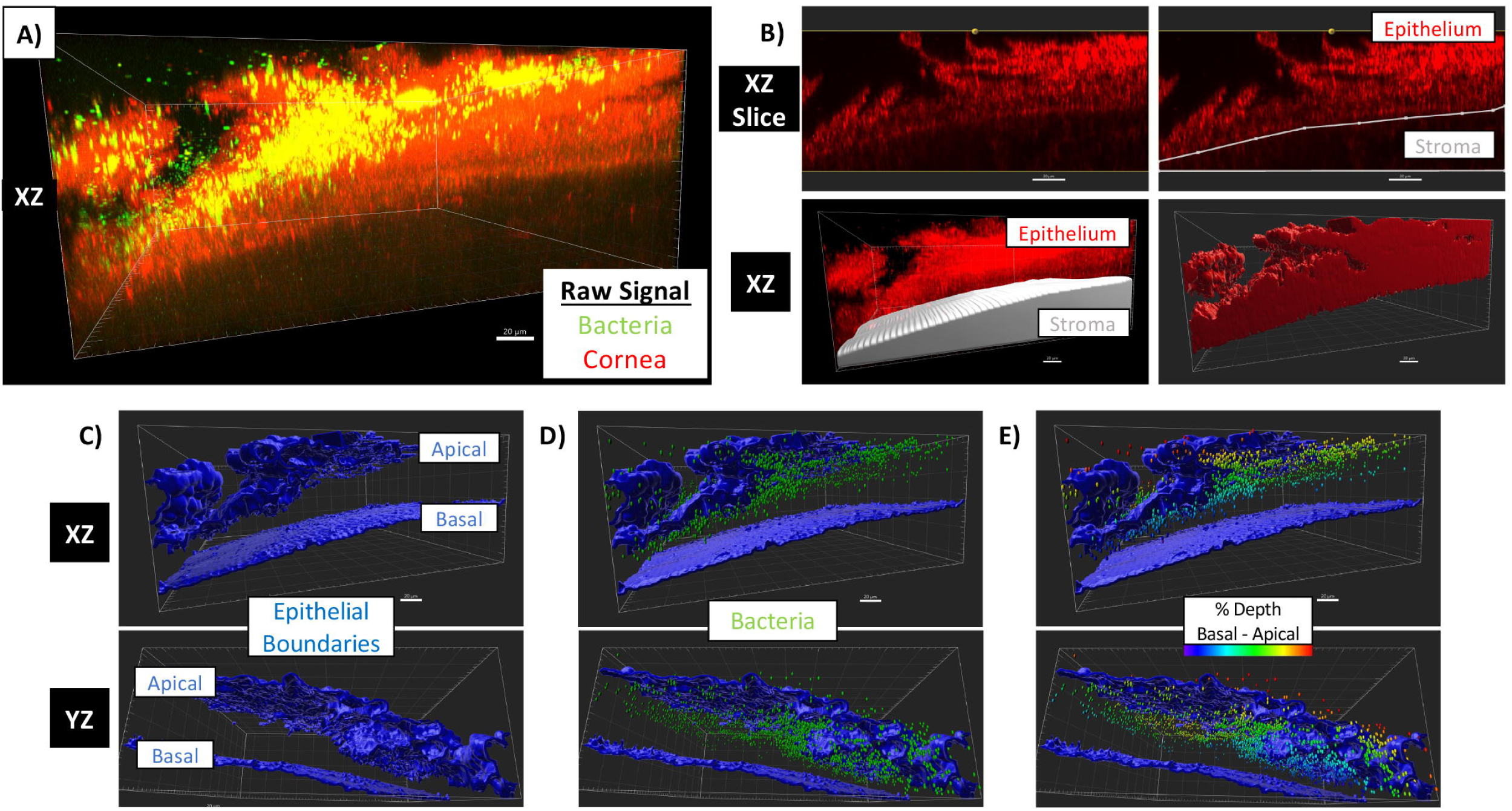
Method for quantitative imaging and analysis of bacterial traversal of the corneal epithelium. Whole murine eyeballs *ex vivo* were blotted with tissue paper, incubated in 0.1M EGTA solution (1 h), then infected with ∼1 x 10^11^ CFU/ml *P. aeruginosa* expressing GFP. Eyes were fixed, whole mounted *en face*, and imaged by confocal microscopy. (A) Raw image imported into Imaris 9.9 for 3D analysis, confocal reflectance signal shown in red and bacterial fluorescence in green. (B) Boundary between stroma and epithelium manually delineated in sequential XZ slices to exclude signal from stroma and create epithelium object. (C) Objects representing apical and basal boundaries of epithelium. (D) Bacteria automatically identified by fluorescent signal quality. (E) Bacteria colored by their depth of traversal, measured as a percentage of its position from the apical to basal boundaries.

**Figure 2.**
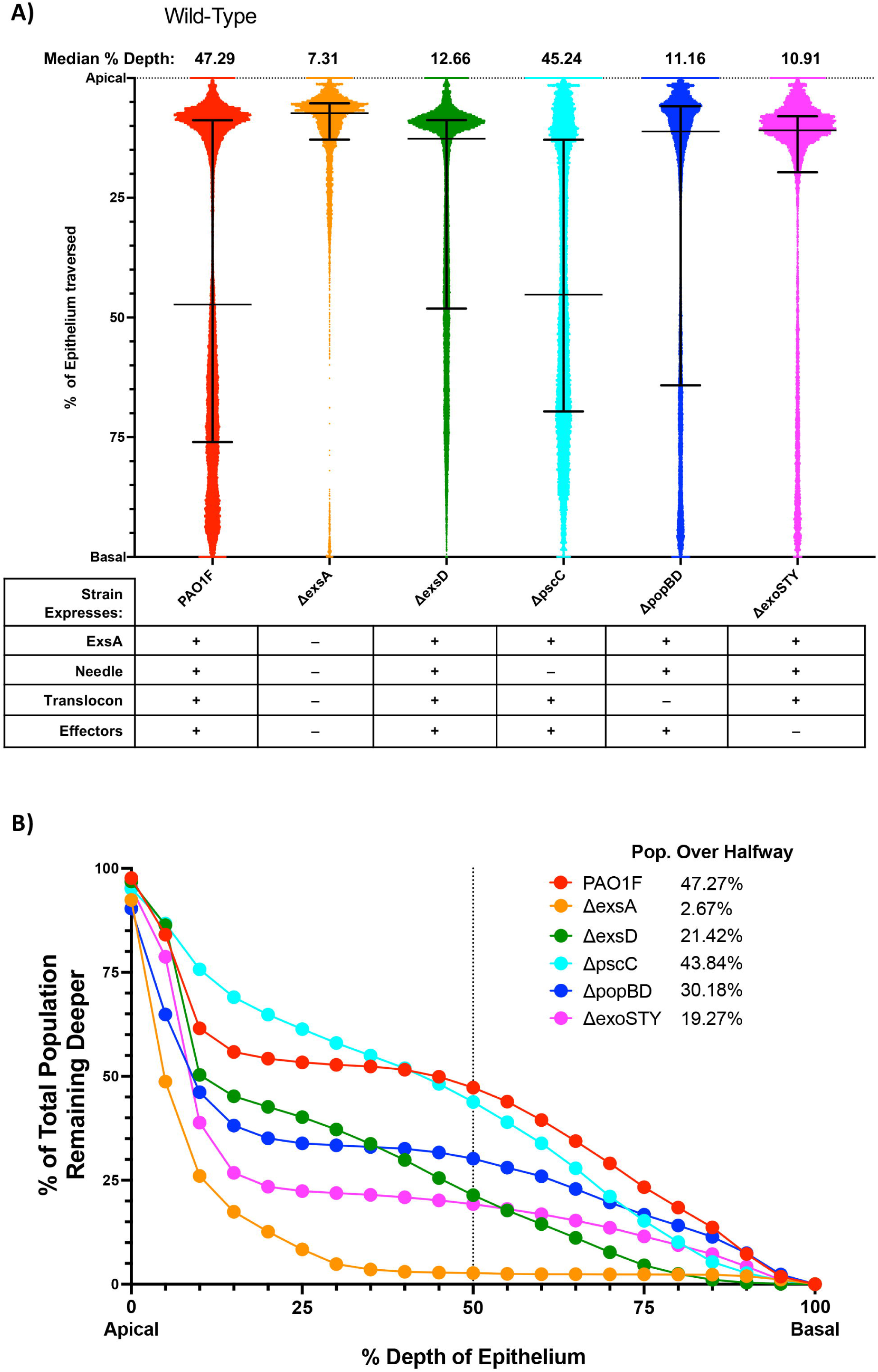
Epithelial traversal by wild-type or T3SS mutants. (A) Traversal depth after *ex vivo* infection by PAO1F and T3SS mutants in genes encoding the transcriptional activator *exsA*, transcriptional repressor *exsD*, known exotoxins *exoSTY*, needle component *pscC*, or translocon pore *popBD*. Each data point represents the relative position (% depth) of one bacterium between the apical and basal boundaries of the 3D corneal epithelium. All data points pooled from ≥ 4 eyes per group. Error bars show the median +/- interquartile range. Note: All comparisons between the T3SS mutants with the Δ*exsA* mutant were significant as were all comparisons with wild-type PAO1F (P < 0.0001, Kruskal-Wallis test with Dunn’s multiple comparisons). (B) Cumulative histogram graphing traversal data (from Fig. 2A) of wild-type and T3SS mutants as a percent of the total bacterial population found deeper in the epithelium. The vertical line indicates halfway through the epithelium, highlighting the proportion of each strain that traversed more than 50% of the epithelial layer.

The first experiment was conducted using *P. aeruginosa* strain PAO1F wild-type compared to an isogenic Δ*exsA* mutant: corneas were incubated with 600 μl of ∼10^11^ CFU/ml bacteria for 6 h (see Methods). The goal was to compare the outcome to our prior study that used more rudimentary methods. For wild-type, the median traversal depth of individual bacteria was 47.29% of the epithelial thickness, the upper quartile reaching at least 76.02% depth (Fig. 2A). Fig. 2B shows that 47.27% of the bacterial population penetrated beyond 50% thickness of the epithelium. In contrast, median penetration depth for Δ*exsA* mutant was only 7.31% depth, the upper quartile being at only 12.86% depth (Fig. 2A). Only 2.67% penetrated beyond the midway point (50% depth) (Fig. 2B). While these findings confirmed the importance of the T3SS in corneal epithelial traversal by *P. aeruginosa* shown in our earlier publications, they provided more granularity on how it impacts bacterial location/distribution. The pattern that emerged showed that the *exsA* mutants lacking the entire T3SS were not completely defective in ability to penetrate beyond the epithelial surface, with significant variability among the population. While wild-type penetrated more deeply than T3SS mutants over the 6 h time span of these experiments, there was again significant variability with only a fraction fully traversing to the level of the underlying basal lamina. For this reason, the capacity to penetrate the corneal epithelium is referred to as <u>traversal efficiency</u> for the remainder of this manuscript.

### Efficient traversal involves T3SS effectors and the T3SS translocon but does not require the T3SS needle apparatus

To explore which ExsA-regulated T3SS components are required for traversal efficiency, we compared wild-type *P. aeruginosa* to mutants lacking the T3SS needle apparatus (Δ*pscC*), the T3SS effectors (Δ*exoSTY*), or the T3SS translocon proteins (Δ*popBD*).

As shown in Fig. 2A, both the T3SS effector mutants (Δ*exoSTY*) and the T3SS translocon mutants (Δ*popBD*) were defective in traversal efficiency compared to wild-type reaching median traversal depths of 11.16% and 10.91% respectively, comparable to the Δ*exsA* mutant. Compared to the Δ*exsA* mutant, a greater proportion of the Δ*popBD* or Δ*exoSTY* mutant populations traversed to over 50% depth; 30.18% and 19.27% respectively (Fig. 2B). Surprisingly, the Δ*pscC* mutant lacking the T3SS needle, was the most traversal efficient of all of the mutants examined, with median traversal at 45.24% depth, similar to wild-type.

These outcomes were surprising given that the T3SS needle is considered important for T3SS functions. Thus, we additionally tested a Δ*exsD* mutant that is constitutively active for the T3SS and therefore consistently expresses T3SS components (including the needle). This differs from wild-type which instead requires induction by host cell contact or low calcium to express the T3SS, and it is effectively the opposite of a Δ*exsA* mutant which consistently lacks T3SS expression. Results with the Δ*exsD* mutant showed that constitutive expression of the T3SS actually <u>interferes</u> with traversal efficiency, showing a median traversal depth of only 12.66% (Fig. 2A), with only 21.42% of the population penetrating beyond 50% depth (Fig. 2B). Representative examples of epithelium traversal by wild-type bacteria and T3SS mutants are shown in Supplemental Fig. S1.

Since traversal efficiency data within these different populations was not normally distributed, the Kruskal-Wallis test with Dunn’s multiple comparisons was used to compare each mutant to the Δ*exsA* mutant. Due to the very large sample sizes (10,000-60,000 datapoints each group), all comparisons were found to be highly statistically significant (P < 0.0001), even when median values and distributions appeared visually comparable (Fig. 2A). This was also true when comparing mutants to wild-type. For this reason, the magnitude and the direction of the differences are important to consider in evaluating the biological significance of the outcomes. Taking those parameters into consideration, the data showed that traversal efficiency was supported by combined efforts of the T3SS effectors and the T3SS translocon. They further suggested that the T3SS needle was not required, and that constitutively expressing it and other T3SS factors in the entire population actually <u>detracted</u> from traversal efficiency. Since this result was unexpected, we used Sanger sequencing to re-confirm that the Δ*pscC* mutant used contained a clean deletion of *pscC* (data not shown). In other controls, the T3SS-GFP reporter pJNE05 (P*_exoS_* expression) was used to confirm that P*_exoS_* was activated upon EGTA-exposure similarly to wild-type in each of the mutants shown to be defective in traversal in Fig. 2: Δ*popBD*, Δ*exoSTY* and Δ*exsD* (Supplemental Fig. S2). Since P*_exoS_* expression is a reliable surrogate for *exsA* expression, this outcome confirmed that the mutants were not defective for T3SS induction, at least under the conditions used.

### Complementation of ExsA in the Δ*pscU-L* T3SS operon mutant restored T3SS effector-dependent traversal efficiency

The above results using T3SS needle mutants and constitutive expression of the T3SS showed that the capacity to make the T3SS needle did not correlate with efficient traversal despite involvement of the T3SS effectors. Here, we directly tested the hypothesis that the effectors can function without the needle.

First, we generated a mutant lacking the 36 genes in the main T3SS operons, from Δ*pscU* to Δ*pscL* (designated Δ*pscU-L*). This mutant still encodes the known T3SS effectors in PAO1 (ExoS, ExoT and ExoY) and their chaperones under endogenous promoters. Results showed the Δ*pscU-L* mutant had low traversal efficiency (Fig. 3A, B) thereby phenocopying the Δ*exsA* mutant. This was to be expected since the Δ*pscU-L* mutant lacks the gene encoding ExsA in addition to genes encoding all of the T3SS machinery proteins, and while it does encode the T3SS effectors, these are not transcribed, translated or released without ExsA.

**Figure 3.**
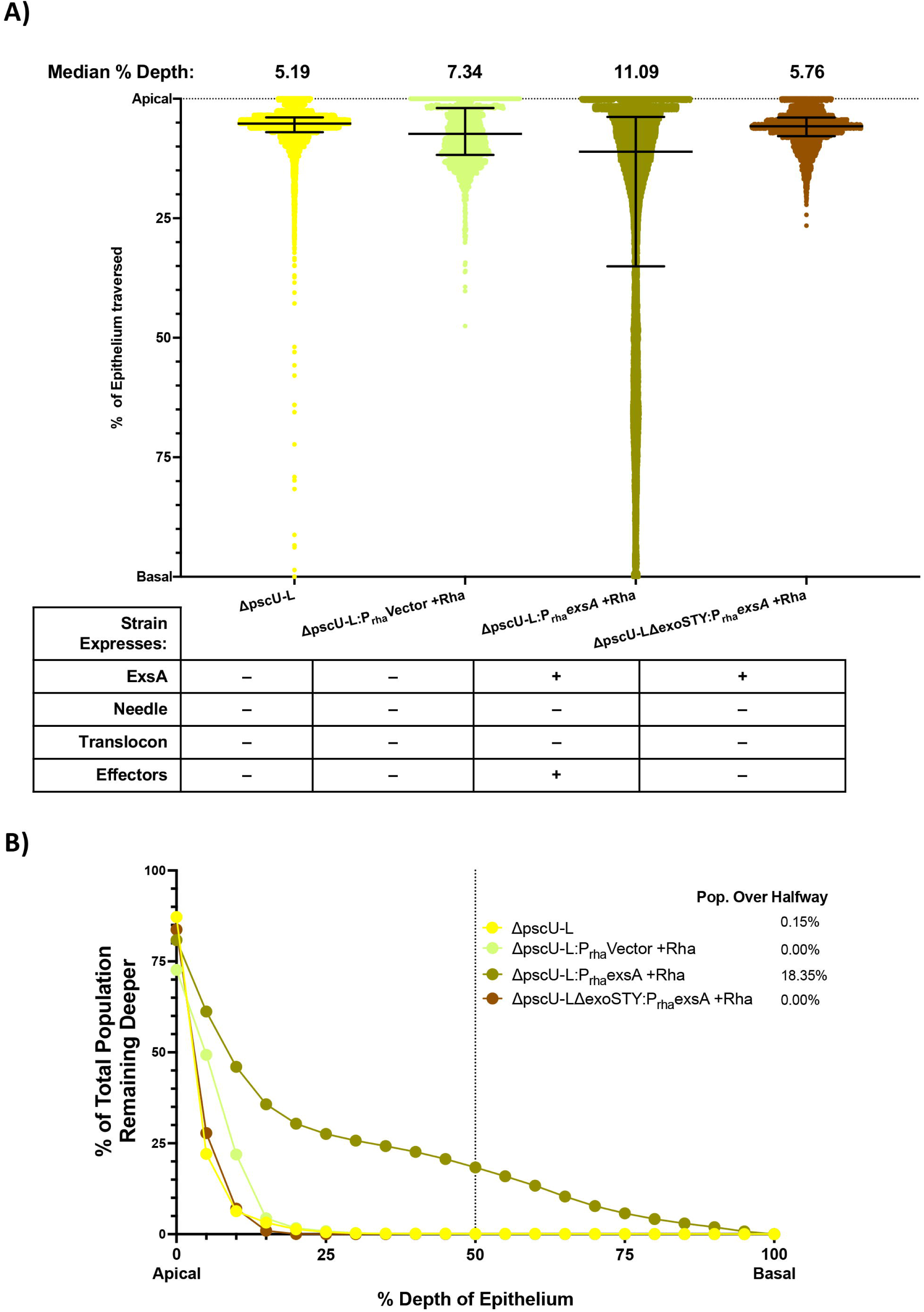
ExsA complementation in a T3SS operon-deficient background. (A) Traversal depth after *ex vivo* infection by a Δ*pscU-L* mutant which lacks all T3SS genes except *exoSTY* (and their respective chaperones) and a Δ*pscU-L*Δ*exoSTY* mutant which also lacks effector toxins. Controls include expression of an empty vector or *exsA* induced by overnight growth on and inclusion of 2% rhamnose during infection. Data were pooled from ≥ 3 eyes per strain. Error bars show median +/- interquartile range. All comparisons between groups were significant at P < 0.0001 except for Δ*pscU-L* vs. Δ*pscU-L*Δ*exoSTY*:P_rha_-*exsA* + rhamnose, which had P = 0.025 (Kruskal-Wallis test with Dunn’s multiple comparisons). (B) Cumulative histogram graphing traversal data (from Fig. 3A) of T3SS operon-deficient mutants with ExsA complementation or controls as a percent of the total bacterial population found deeper in the epithelium. The vertical line indicates halfway through the epithelium, highlighting the proportion of each strain that traversed more than 50% of the epithelial layer.

Next, we tested if expressing the T3SS effectors in this mutant lacking T3SS machinery related proteins could rescue traversal efficiency. This was done by complementing the mutant with ExsA using a chromosome-integrating vector to induce expression of *exsA* under a rhamnose-sensitive promoter. This vector was first tested in the background of a Δ*exsA* mutant using a GFP reporter pJNE05 for *P_exoS_* expression as a surrogate for *exsA* induction (as above, Supplemental Fig. S2). Rhamnose addition induced GFP expression and it significantly increased bacterial traversal compared to the no rhamnose control for the Δ*exsA*:P_rha_*exsA* mutant (Supplemental Fig. S3A, B). We next showed that Rhamnose-induction of *exsA* expression in the Δ*pscU-L* mutant background increased median traversal to 11.09% depth versus the rhamnose-induced vector control at 7.34% depth (Fig. 3A), and it also increased the percentage of the population that traversed to at least 50% depth to 18.35% vs. 0.00 % (Fig. 3B). Showing that partial restoration of traversal efficiency by Δ*pscU-L*:P_rha_*exsA* depended on the T3SS effectors, it was abrogated when the three exotoxin genes were also deleted (Median: 5.76% depth, population over halfway 0.00%) (Fig. 3A, B). An *in vitro* control with pJNE05 confirmed activation of P*_exoS_* in the Δ*pscU-L* mutant after *exsA* complementation with rhamnose addition relative to a vector-complemented control: Mean +/- SD P*_exoS_* expression (FITC/OD_600_) for Δ*pscU-L*:P_rha_*exsA* + rhamnose was 3397 ± 1033 relative fluorescence units versus 1567 ± 1235 for Δ*pscU-L*:P_rha_Vector + rhamnose (P = 0.0061, Student’s t-Test).

In Fig. 3A all comparisons between groups were significant (P < 0.0001 for most comparisons, and P = 0.025 comparing Δ*pscU-L* [no rhamnose] *vs.* Δ*pscU-L*Δ*exoSTY*:P_rha_*exsA* [+ rhamnose], Kruskal-Wallis test with Dunn’s multiple comparisons). The latter comparison compared two mutants both unable to express T3SS effectors or any of the T3SS machinery-related proteins, only one able to express the transcriptional activator ExsA (Fig. 3A, B). Thus, ExsA expression in the absence of those other components had the least impact on traversal efficiency. Taken together, these results show that ExsA-driven expression of *exoSTY* can increase traversal efficiency of a mutant lacking all the T3SS machinery and thus one or more effectors are necessary for traversal. However, it remains possible that ExsA regulates the expression of a factor(s) outside of the main T3SS operon that is/are needed for effector-mediated traversal.

### The T3SS translocon contributes to traversal efficiency beyond roles of the T3SS effectors, also independently of the T3SS needle

To explore if the T3SS translocon contributes to traversal efficiency beyond the impact of the T3SS effectors, we studied deletion of *popB* in the background of a Δ*exoSTY* mutant (already reduced in traversal efficiency compared to wild-type). This led to a small but statistically significant further reduction in median traversal depth: Δ*exoSTY* median at 10.91% depth vs. Δ*popB*Δ*exoSTY* median at 8.10% depth (Fig. 4A, Δ*exoSTY* data reproduced from Fig. 2) (P < 0.0001, Kruskal-Wallis test with Dunn’s multiple comparisons). A much bigger reduction was seen in the percent of population traversing beyond the 50% point: Δ*exoSTY* 19.27% vs. Δ*popB*Δ*exoSTY* 1.62%.

**Figure 4.**
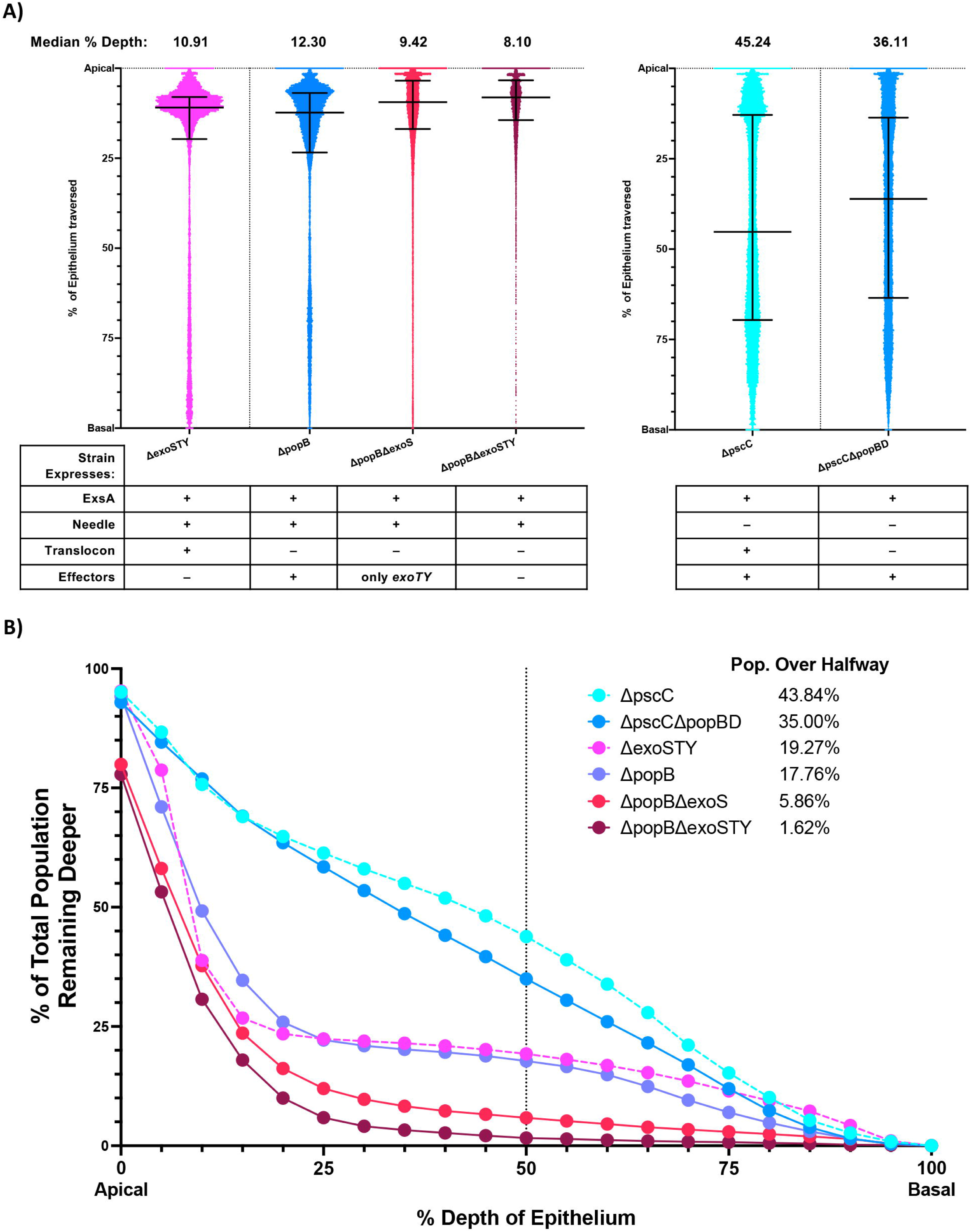
Traversal by translocon and needle- or exotoxin-deficient mutants. (A) Traversal depth after *ex vivo* infection by T3SS translocon mutants in a background of Δ*pscC*, Δ*exoS*, or Δ*exoSTY*. Data from Fig. 2A was used for Δ*pscC* and Δ*exoSTY* mutants. Data pooled from ≥3 eyes per strain. Error bars show median +/- interquartile range. All comparisons between groups were significant (P < 0.0001, Kruskal-Wallis test with Dunn’s multiple comparisons) except for Δ*exoSTY* vs. Δ*popB.* (B) Cumulative histogram graphing traversal data (from Fig. 4A) of the T3SS mutants as a percent of the total bacterial population found deeper in the epithelium. The vertical line indicates halfway through the epithelium, highlighting the proportion of each strain that traversed more than 50% of the epithelial layer.

We next tried the opposite experiment, deleting a T3SS effector gene (*exoS*) in a T3SS translocon mutant (*popB*). This also showed separable contributions for the T3SS effectors and the T3SS translocon, the median depth for a Δ*popB* mutant at 12.30% depth vs. median for Δ*popB*Δ*exoS* mutants at 9.42% depth (Fig. 4A) (P < 0.0001, Kruskal-Wallis test with Dunn’s multiple comparisons). Again, the difference became more obvious when assessing the percent of the population penetrating beyond 50% depth; Δ*popB* 17.76% vs. Δ*popB*Δ*exoS* 5.86% (Fig. 4B). Taken together, these results showed that T3SS translocon proteins and T3SS effector proteins can have additive impacts on traversal efficiency and specifically implicate ExoS and PopB. However, difference in traversal beyond 50% depth shown between Δ*popB*Δ*exoS* 5.86% and Δ*popB*Δ*exoSTY* 1.62% implicate a role(s) for ExoT and/or ExoY.

Having already shown that the T3SS needle was dispensable for effector mediated traversal, we next asked if it was needed for the translocon’s contribution. Thus, we mutated the translocon pore proteins PopB and PopD (Δ*popBD* mutant) in the background of the T3SS needle mutant (Δ*pscC*) already shown to be traversal competent (Fig. 2). This reduced traversal efficiency of the needle mutant: Δ*pscC* mutant median 45.24% depth, population over halfway 43.84% vs. Δ*pscC*Δ*popBD* mutant median at 36.11% depth, population over halfway at 35.00% (Fig. 4A, B, Δ*pscC* data reproduced from Fig. 2) (P < 0.0001, Kruskal-Wallis test with Dunn’s multiple comparisons). This showed that the T3SS translocon can contribute to traversal efficiency in the absence of the T3SS needle.

All traversal results and strain characteristics are summarized in Table 1. Together, these outcomes implicate two separate contributors to T3SS-mediated traversal efficiency: the T3SS effectors (including ExoS), and the T3SS translocon pore (including PopB), each functioning independently of the T3SS needle which can instead play an inhibitory role. Interestingly, the <u>magnitude</u> of their individual contributions appear similar in magnitude, the Δ*exoSTY* and Δ*popB* mutants showing no significant difference in impact (Fig. 4).

**Table 1.**
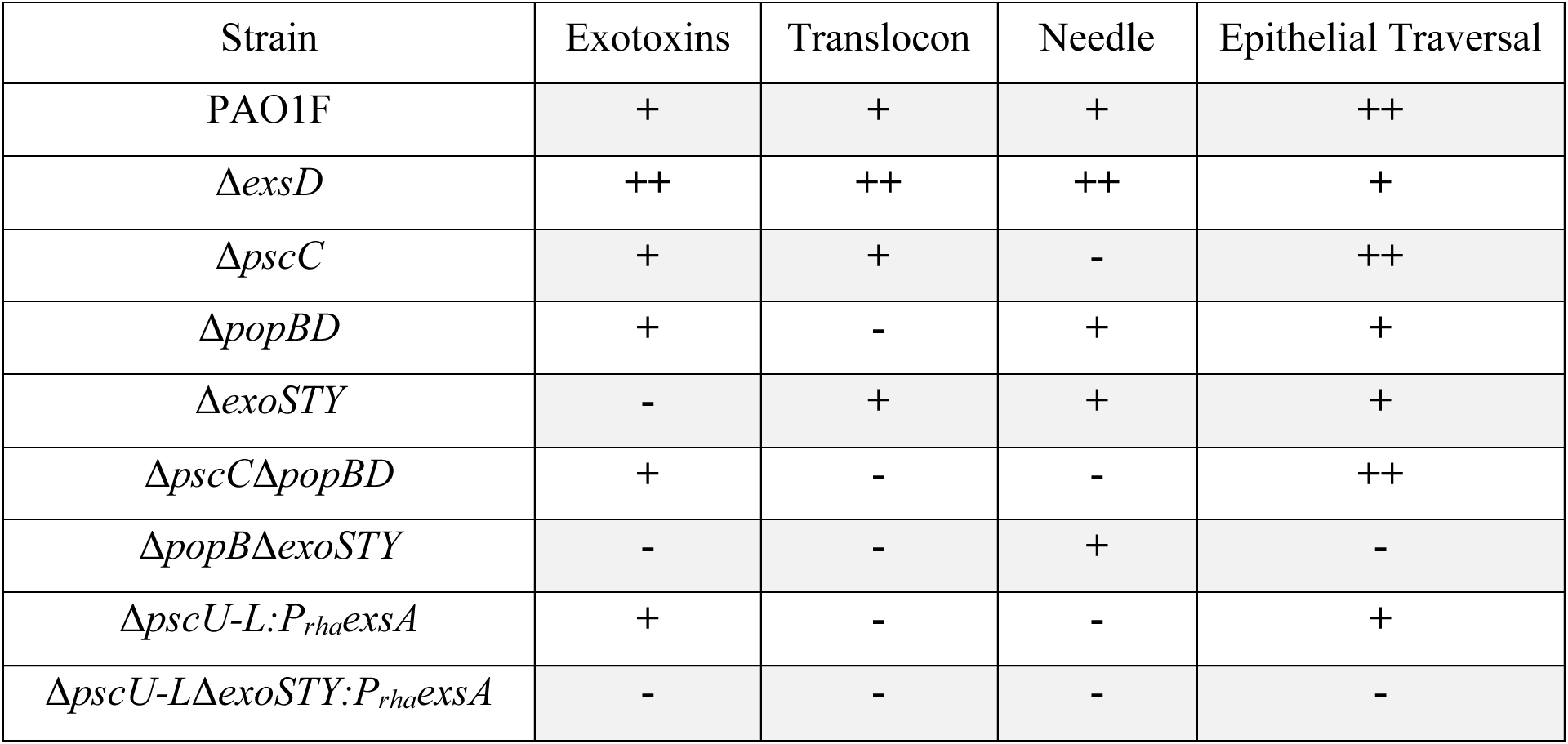
Properties and epithelial traversal results for *P. aeruginosa PAO1F and T3SS mutants*.

### Exotoxin S protein can be released by a T3SS operon mutant

Results in Figs. 2-4 show that the T3SS effectors can contribute to traversal efficiency without the T3SS needle genes. Our current understanding of T3SS transcription and expression is that in the presence of Ca^2+^ or absence of host cell contact, low levels of T3SS expression occur due to repression of ExsA by ExsD (63). Upon host cell contact or Ca^2+^ deprivation, the T3SS regulator ExsE is secreted, leading to ExsA de-repression and T3SS transcription. *In vitro*, Δ*pscC* needle mutants do not secrete ExsE and do not activate high levels of T3SS expression (64). To test if an alternate mechanism of T3SS effector toxin expression might be operating in our *ex vivo* traversal model or T3SS mutants, the Δ*exsA* or Δ*pscU-L* mutant carrying rhamnose-inducible *exsA* were tested for ExoS protein expression *in vitro* using LB broth without EGTA calcium chelation to specifically test rhamnose induction compared to controls (vector only or no rhamnose). After growth in broth culture to ‘mid-log’ phase, bacterial pellet and supernatant samples were collected and probed for ExoS *via* affinity-purified polyclonal antibody using SDS-PAGE and Western immunoblot as described previously (64). ExoS protein was detected in the pellet and supernatant of Δ*exsA*:P_rha_*exsA* with rhamnose induction (as expected) and in Δ*pscU-L*:P_rha_*exsA* with inclusion of rhamnose but not in its vector control (Fig. 5). These data confirmed that ExsA-driven ExoS expression and release can occur for mutants lacking needle protein genes.

**Figure 5.**
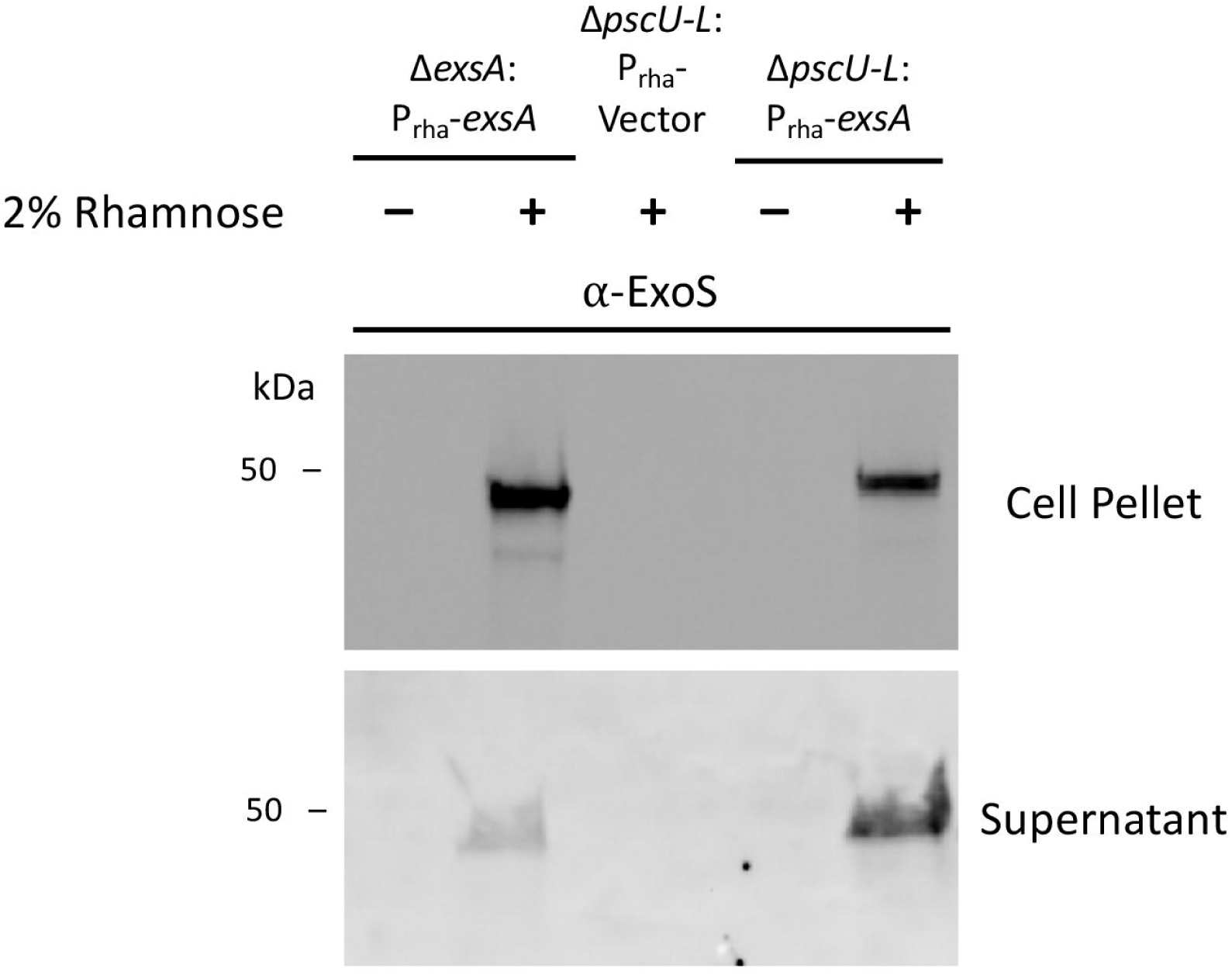
Rhamnose-induction of ExsA-driven ExoS expression in both bacterial cells and supernatant of a needle-deficient mutant Δ*pscU-L*. Western immunoblot showing bacterial cell pellets and supernatant fractions collected from Δ*exsA*:P_rha_*exsA*, Δ*pscU-L*:P_rha_Vector, Δ*pscU-L*:P_rha_*exsA* after growth to ‘mid-log’ phase in LB media, with induction of *exsA* by 2% rhamnose inclusion for the conditions indicated. The location of bands corresponding with ExoS (∼49 kDa) detected by affinity-purified antibody and chemiluminescence are evident below the protein standard annotated at 50 kDa.

## Discussion

Previously, we showed that the transcriptional activator of the T3SS (ExsA) is required for *P. aeruginosa* to traverse susceptible multilayered epithelium in the context of live tissue, shown using the eyes of immunocompetent mice (38, 62). Here, we explored the role of T3SS components.

The cornea has several advantages over other tissues for this type of investigation being readily accessible for manipulation or imaging, optically clear when healthy allowing imaging without 2-photon microscopy or tissue clearing protocols and is relatively separated from other body sites reducing complexity associated with cross-talk that can occur during infection. Its epithelial surface also lacks a microbiome (35, 36), allowing a pathogen to be studied alone or in combination with other bacteria in a controlled fashion. Use of subtle epithelial injury to enable susceptibility to bacterial traversal, as done in this study using tissue paper blotting then EGTA treatment, preserves the underlying basal lamina barrier, prohibiting bacterial access to the vulnerable stroma and allowing study of epithelial traversal by bacteria without complexities introduced by later steps of infection development. Thus, this mouse eye model for studying epithelial traversal shares some advantages of *in vitro* cell/tissue culture models in reducing complexity but benefits from *in situ* context.

A challenge of *in situ* tissue imaging for quantitively localizing bacteria is variations in tissue shape that can make it difficult to ascertain position relative to upper and lower boundaries across an entire tissue sample. Here, we developed methods that account for the cornea’s curved surface, variations in thickness and topography in different regions and resulting from cell exfoliation, and any changes to these parameters after exposure to bacteria. This allowed for more precise quantitation of bacterial location between the epithelial surface and underlying basal lamina, separately accounting for all individual bacteria present in the epithelial layer.

To calibrate outcomes to our previously published methods, we first compared wild-type *P. aeruginosa* to mutants lacking the entire T3SS *(exsA* mutants) which confirmed the critical role for the T3SS in corneal epithelium traversal. However, the more accurate localization method showed traversal was not an absolute quality, with both wild-type *P. aeruginosa* and *exsA* mutants showing a distribution/spread for individual bacteria in their penetration rates. Thus, we referred to “traversal efficiency” rather than “traversal” to describe subsequent outcomes.

When roles of specific T3SS components were examined, some results aligned with *in vitro* cell culture study findings (65). For example, mutants lacking genes for known T3SS exotoxins had reduced traversal efficacy compared to wild-type. However, our study showed that mutants in T3SS translocon pore proteins also showed reduced traversal efficiency, with mutants lacking both translocon and exotoxin proteins showing an even more profound defect similar to *exsA* mutants unable to express any T3SS components. This implicated both the T3SS effectors and the T3SS translocon proteins, their roles being additive.

Not all outcomes aligned with *in vitro* study findings. Mutants lacking the T3SS needle were just as traversal efficient as wild-type *in situ.* This was a surprising and confusing result because the T3SS needle is generally thought critical for T3SS function, including for secretion or expression of the very same T3SS components we found required for traversal efficiency. Thus, we used a mutant lacking the entire T3SS operon except the genes encoding the effector exotoxins (Δ*pscU-L* mutant) (66) and induced effector expression by complemented it with rhamnose inducible *exsA* (the transcriptional activator of all T3SS-related genes). This promoted traversal efficiency in the mutant despite it lacking T3SS needle and other machinery related proteins. Confirming it was T3SS effector gene expression driving the phenotype in the T3SS operon mutant not than other genes outside the T3SS operon, traversal was no longer promoted by *exsA* complementation when the effector genes were also mutated (i.e. Δ*pscU-L exoSTY* mutant). Since Δ*pscU-L* mutants lack T3SS translocon genes additionally needed for full traversal efficiency, it was not surprising that *exsA* complementation did not fully restore the operon mutant to wild-type traversal efficiency. Indeed, efficiency approximated translocon (*popBD*) mutants that similarly encode T3SS effectors but not translocon proteins, but that differ in encoding T3SS needle proteins. Together these results confirmed that the T3SS needle protein genes are not needed for T3SS effector-mediated traversal efficiency in this *in situ* model.

Since T3SS needle proteins are thought necessary for exporting T3SS effectors out of bacteria and for injecting them into host cells this data raised two related questions. Can T3SS effectors become extracellular without the needle, and if so, how do they impact traversal without being injected into host cells?

Western immunoblotting confirmed the presence of ExoS in both supernatant and pellet of *in vitro* grown Δ*pscU-L*:P_rha_-*exsA* after rhamnose induction, showing that ExoS can become extracellular without needle protein genes. While this might be an unusual feature of this mutant, that would not explain results with other mutants that also show the needle is dispensable for T3SS mediated traversal efficiency. A potential explanation is that bacterial lytic cell death releases effectors, e.g. explosive cell lysis in *P. aeruginosa* (67), and a feature of other toxin release by other bacteria (68, 69) and quite possible given the dense bacterial inoculum and routine ‘natural’ bacterial cell death as a result of nutrient deprivation or from formation of biofilms (67). In the context of our *in situ* assay, the presence of host-derived antimicrobial peptides (29, 33, 70) could further contribute to bacterial cell death. Alternatively, T3SS proteins are packaged into OMVs (Outer Membrane Vesicles) that are released into the extracellular environment, as shown for *Salmonella enterica* T3SS-1 effectors and translocation proteins (71). Indeed, some T3SS proteins were efficiently secreted *via* OMVs by mutants lacking a functional T3SS needle, and effector toxins were shown delivered *via* OMVs into the cytoplasm of epithelial cells where they enhanced pathogenesis of T3SS-deficient mutants (72). Pathogenic *E. coli* (O157:H7) can also efficiently package T3SS translocation and effector proteins into OMVs in the absence of essential T3SS needle proteins (73). Our own work showed that exposing *P. aeruginosa* to ocular surface tear fluid or to purified lysozyme (a tear fluid ingredient) generates OMVs that are cytotoxic to the corneal epithelium of mouse eyes *in vivo*. We also showed that priming the corneal surface with these OMVs reduces defense against bacterial adhesion, and that tear fluid/lysozyme triggered OMVs contain a ∼48 kDa protein similar in size to ExoS (∼49 kDa) and ExoT (∼53 kDa) (74). Another potential pathway for delivering T3SS effectors into a host cell without the T3SS needle could be from an intracellular location. In this regard, it is interesting that the results specifically implicated PopB and ExoS in traversal efficiency as our work has shown critical roles for both in the intracellular lifestyle of *P. aeruginosa* (24–27, 75). Notwithstanding, even T3SS mutants are internalized by epithelial cells and can persist inside intracellular vacuoles that can subsequently fail to contain them (24–26, 75–78). In other words, T3SS factors can facilitate but are not essential for bacterial entry into a host cell’s cytoplasm. Inside epithelial or other cells, bacteria (including *P. aeruginosa*) can use factors beyond T3SS factors to exert impacts on the host cell, and the host cell can detect intracellular bacteria and in response alter its biology and/or induce programmed cell death. Others have shown that extracellular ExoS itself can be internalized by eukaryotic cells, and activates host cell TNF-α responses by triggering surface TLRs (79). While mechanisms might overlap with some of the above, ExoS has also been shown to affect host cell function from an extracellular location (80).

Given these many possibilities, a separate study will be needed to determine how T3SS effector mediated traversal occurs without the T3SS needle. Importantly, much of our understanding of gene expression in *P. aeruginosa* has been done using *in vitro* methods, often without host cells present. Given its many environmental sensors and regulators of gene expression, regulation of the *P. aeruginosa* T3SS *in situ/in vivo* could differ vastly from what has been shown *in vitro*. It is also likely to be context dependent/complex due to microenvironments, and lack of homogeneity and synchrony. Indeed, the result showing that *exsD* mutants (which differ from wild-type in constitutively expressing the T3SS) are equally as traversal defective as mutants lacking the entire T3SS are particularly poignant and highlight the importance of regulation. Further research on this topic would benefit from more *in situ* experimentation to set the stage for well-designed *in vitro* studies that can then tease apart mechanisms for the phenomena shown relevant to pathogenesis *in situ/in vivo*.

The T3SS translocon proteins (PopB/PopD) were found to play roles additional to the contribution of the effectors. This could relate to their ability to form pores in host cell membranes, which can damage a cell or otherwise alter host cell function. Indeed, PopB-PopD complexes in isolation have been shown to function as a pore-forming toxin by allowing K^+^ efflux, resulting in histone H3 modification and host cell subversion (81). A potential mechanism for the additive impacts of translocon and effector proteins could be pore formation providing an entry mechanism for effector delivery into the host cell. Whether the T3SS needle is also dispensable for translocon-mediated traversal efficiency remains an open question that will require further investigation, although it is suggested by the fact that needle mutants remain fully efficient.

Another factor to consider in interpreting the data is that T3SS needle proteins of *P. aeruginosa* and other Gram-negative bacteria are recognized by mouse (and human) cells causing inflammasome activation (82, 83), which can help the cornea and other tissues recognize, respond and clear *P. aeruginosa* during experimental challenge *in situ* (35, 84). In this way, expressing T3SS needle proteins could be detrimental to bacteria *in situ* (even if not *in vitro*) and using needle-independent mechanism for traversal could be advantageous. Other *in situ* factors (absent *in vitro*) may also impact bacterial viability or growth during traversal and thus data interpretation.

Providing additional insights into pathogenesis, traversing bacteria tended to be bimodally distributed across the epithelium. Typically, there was a cluster in the apical region from 0-15% traversed, few across the midway point, then a group from 50-80% traversed. This distribution may reflect bistability of T3SS expression in *P. aeruginosa* populations (26, 85), and/or a host defense barrier, possibly related to specialized junctions in the suprabasal region of the corneal epithelium that have been shown to be a barrier to leukocytes (86). Also worth noting, the very large sample sizes (10,000-60,000 datapoints for each group) provided such a powerful analysis, that most comparisons yielded statistically significant differences even when they were small. Thus, we considered the magnitude of the differences in central tendency (median) and data spread (upper and lower quartiles) to draw conclusions about biological significance.

In conclusion, this study used novel and powerful image analysis tools and an *in situ* model with pared-down complexity to focus on an early step in pathogenesis, traversal of a superficial multilayered epithelium. The goal was to determine which T3SS components contribute. Results showed roles for T3SS effectors, T3SS translocon proteins, and regulation of T3SS expression. Differing from *in vitro* culture studies and challenging our general understanding about how the *P. aeruginosa* T3SS supports infection pathogenesis, the T3SS needle was dispensable. Follow up experiments confirmed that the T3SS effector ExoS could become extracellular in the absence of the T3SS needle. How T3SS effectors and translocon proteins influence host cells to promote traversal without the T3SS needle will require further work. Published knowledge about *P. aeruginosa* and other Gram-negative bacteria suggest multiple possibilities (67, 72, 73, 87).

Outcomes of this study may or may not be applicable to how *P. aeruginosa* traverses epithelial barriers other than the mouse corneal epithelium, but if so they might generally inform efforts to develop therapies targeting the T3SS (88, 89). Importantly, the results highlight the need to move beyond *in vitro* and cell culture studies when studying pathogenesis. At the same time, they support complementary *in vitro* and cell culture experimentation that can further reduce complexity when teasing apart details, and to study regulation in a more controlled system.

The model and imaging methods we have developed enable a single early stage of infection pathogenesis to be studied *in situ* in the absence of various confounders generally present in infection models (e.g. inflammation), and they enable individual bacteria to be visualized across the entire tissue with their location quantified. These methods could prove useful for studying pathogenic strategies of other microbes also able to traverse epithelial layers during infection, including those not considered to be eye pathogens.

## Materials and Methods

### Bacteria

*Pseudomonas aeruginosa* strain PAO1F was used for all experiments. Mutants in T3SS components were either obtained from in-house strain collections or generated using two-step allelic exchange (90). For this, pEXG2 vectors containing overlapping upstream and downstream regions of the genes of interest (excluding the ORF) were cloned and transformed to *E. coli* SM10 (λ*pir*), which were then used as donor strains to deliver the vector to recipient *P. aeruginosa* strains. Successful merodiploid colonies underwent sucrose counter-selection and PCR verification of mutagenesis. For integration of rhamnose-inducible ExsA, Dr. Arne Rietsch provided strains generated by Tn7 vector-mediated insertion and Flp-mediated excision of antibiotic resistance markers (91). All bacteria, mutants and plasmids used are shown in Table 2. In control experiments not shown, T3SS mutants did not show any growth defects relative to PAO1 in trypticase soy broth culture at 37 °C *in vitro*.

**Table 2.** Strains, mutants, and plasmids used in this study.

| Strain or mutant | Description | Source |
| --- | --- | --- |
| PAO1F | Wild-type *P. aeruginosa*; All mutants made within this background | Dr. Alain Filloux *via* Dr. Arne Rietsch (94) |
| $\Delta exsA$ | *exsA* mutant; Constitutively T3SS<sup>Off</sup> | Dr. Arne Rietsch |
| $\Delta exsD$ | *exsD* mutant; Constitutively T3SS<sup>On</sup> | Dr. Timothy Yahr (63) |
| $\Delta pscC$ | *pscC* mutant; Lacking needle | Dr. Arne Rietsch (95) |
| $\Delta popBD$ | *popBD* double mutant; Lacking translocon pore | Dr. Arne Rietsch (95) |
| $\Delta exoSTY$ | *exoSTY* triple mutant; Lacking known exotoxins | Dr. Arne Rietsch (95) |
| $\Delta pscU-L$ | Total T3SS-operon mutant (knockout of all genes *pscU* to *pscL*); called $\Delta U-L$ | Dr. Arne Rietsch (66) |
| $\Delta exsA:Prha-A$ | *exsA* mutant with chromosome-integrated rhamnose-inducible *exsA* | Dr. Abby Kroken (27) |
| $\Delta pscU-L:Prha-A$ | $\Delta U-L$ with rhamnose-inducible *exsA* | This study |
| $\Delta pscU-L:Prha-V$ | $\Delta U-L$ with rhamnose-inducible vector control | This study |
| $\Delta pscU-L$<br>$\Delta exoSTY:Prha-A$ | $\Delta U-L$ lacking all known exotoxins, with rhamnose-inducible *exsA* | This study |
| $\Delta popB\Delta exoSTY$ | *popB* mutant lacking all known exotoxins | Dr. Victoria Hritonenko (75) |
| $\Delta pscC\Delta popBD$ | *popBD* mutant lacking T3SS needle | This study |
| $\Delta popB\Delta exoS$ | *popB* mutant lacking *exoS* | This study |
| *E. coli* SM10($\lambda pir$) | Mutagenesis vector donor strain | New England Biolabs |
| Plasmid | Description (Resistance conferred, $\mu g/ml$) | Source |
| pSMC2 | Constitutive GFP (Carbenicillin, 400) | (96) |
| pJNE05 | T3SS-GFP (*PexoS*). Plasmid encoding *gfp* fused to the ExsA-dependent promoter of *exoS*. (Gentamicin, 200) | (97) |
| pEXG2-$\Delta pscU-L$ | Integrating suicide plasmid for mutagenesis | Dr. Arne Rietsch |
| pEXG2-$\Delta exoS$ | Integrating suicide plasmid for mutagenesis | Dr. Arne Rietsch |
| pUC18-mini-Tn7T-PrhaB-*exsA* | Delivery vector for integrating *exsA* under Rhamnose-sensitive promoter to *attTn7* site (Carbenicillin, 100) | This study |
| pUC18-mini-Tn7T-PrhaB-Vector | Delivery vector for integrating empty expression vector under rhamnose-sensitive promoter to *attTn7* site (Carbenicillin, 100) | This study |
| pTNS1 | Helper plasmid for conjugation of Tn7 integration plasmid (Ampicillin, 100) | (91) |
| pRK2013 | Helper plasmid for conjugation of Tn7 integration plasmid (Kanamycin, 35) | (91) |
| pFLP2 | Sucrose-sensitive vector to excise antibiotic resistance marker from *attTn7* site | (91) |

For all experiments, bacteria were streaked from glycerol stocks stored at -80 °C, plated onto trypticase soy agar (TSA) plates with appropriate antibiotics (see Table 2) and grown at 37 °C overnight. For the rhamnose-induction traversal experiments, bacteria were grown on TSA also supplemented with rhamnose (2 %). For T3SS-induction growth curves with plasmid pJNE05, bacteria were grown in trypticase soy broth (TSB) supplemented with gentamicin (200 μg/ml), monosodium glutamate (100 mM), glycerol (1%) and EGTA (2 mM) or alternatively with rhamnose (2 %) instead of EGTA as specified.

For the *ex vivo* murine model of traversal, one bacterial “lawn-covered” TSA plate was used per bacterial strain per condition. After overnight growth at 37 °C, bacteria were collected with a sterile loop and carefully suspended into serum-free DMEM (Dulbecco’s Modified Eagle medium; Gibco). DMEM was supplemented with 2% rhamnose when indicated. OD_600_ was measured, with A_600_ 1.0 ∼ 4 x 10^8^ CFU/ml used for calculating density. Suspensions of 1.0 x 10^11^ CFU/ml were used for *ex vivo* incubations, and concentrations confirmed by plating serial dilutions for viable counts.

### *Ex vivo* murine model of traversal

All procedures were approved by the Animal Care and Use Committee of the University of California, Berkeley, an AAALAC accredited institution. For all experiments, male or female C57BL6/J or mT/mG mice with tdTomato-labeled cell membranes were used, 6 to 12 weeks old, and 3-6 eyes per experimental condition. Contralateral eyes from the same mouse were used as controls when possible. All experiments were performed *ex vivo*, as mice were euthanized by isoflurane inhalation prior to any further manipulations. After euthanasia, each eye was lightly blotted with a Kimwipe (Kimtech), enucleated, then placed in 0.2µm filter-sterilized 1X Phosphate-Buffered Saline (PBS) in a 48-well dish for subsequent steps. Eyes were rinsed three times in PBS, then incubated in a 0.1M solution of EGTA at room temperature for 1 h. Eyes were rinsed three more times in PBS, then completely submerged in a 600 µl suspension of bacteria, prepared as described above, and incubated at 37 °C, 5 % CO_2_ for 6 h, consistent with our previous studies (62, 92). Following this incubation, eyes were rinsed with PBS three times to remove excess non-adherent bacteria before fixation in 2 % paraformaldehyde (PFA) overnight at 4 °C. Fixed eyes were then whole mounted on a glass cover slip with Loctite superglue with the central cornea facing up before submersion in PBS and imaging.

### Imaging

Whole-mounted fixed corneas were imaged using an Olympus Fluoview FV1000 upright laser scanning confocal microscope with a water immersion 20X objective (NA = 1.0). For each eye, three to five non-intersecting fields of view on the central cornea were selected for imaging with 3x optical zoom (60x total magnification) at 1024 x 1024 resolution and 0.8µm step size. Fluorescent bacteria were imaged using FITC (488/509) and/or RFP (555/580) channels, and corneal cells were imaged using an 640nm excitation channel for confocal reflectance microscopy (CRM) (92, 93).

### Image analysis

Images of corneas were processed on Imaris Software (v 9.9) as follows. First, a “surface object” was manually drawn using the reflectance channel to select the total signal underneath the corneal basement membrane. This object represented the corneal stroma and was used to mask the remaining reflectance signal. This created a new channel with signal from only the epithelium, which was used to automatically generate a new surface object.

From the created Epithelium surface object, a distance transformation was performed, and the basement membrane and apical surface were identified as two new surface objects ∼1µm away from the Epithelium. Variability in the created objects due to differing corneal reflectance intensity was minimized through use of automatic thresholds, and manual surveillance of the entire process ensured consistency in appearances. Next, bacteria were identified through the ‘Spots’ tool, using an XY size of 1.25 µm in the appropriate channel. Parameters for Spots creation were assessed using an automatic threshold, adjusted manually to reduce false positive results (e.g., Spots where there were no bacteria).

For each field of view, the following were measured: total volume and maximum thickness of epithelium, total number of bacteria, and distance of each bacterium from apical surface and basement membrane. Percent depth of traversal for each bacterium was calculated using the formula: % Depth = Distance to Apical Surface / (Distance to Apical + Distance to Basement Membrane) * 100 %. An inverted cumulative histogram was also generated from the % depth for all bacteria in a condition with 5µm bins. The percent of the total population remaining deeper was graphed, so 100 % of a population remained at 0 % traversal depth and 0 % remained at 100 % depth.

### Western immunoblot

Bacteria were inoculated in liquid cultures of LB media (containing 200 mM NaCl, supplemented with 10 mM MgCl_2_) and grown with shaking overnight at 37 °C. Overnight cultures were diluted 1:300 into fresh media, grown for 2 h, then 1 ml of culture was added to 1 ml pre-warmed media with 2 % rhamnose and 1 ml of culture was added to 1 ml pre-warmed media as a control. The bacteria were grown for another hour before the OD_600_ was measured for each tube, which were then placed on ice. 1 ml of each sample was pelleted, and the entire supernatant was added to a fresh tube containing 10 % trichloroacetic acid for precipitation. After 10 min on ice, two washes with acetone were conducted before drying out the precipitate at room temperature. The cell pellet and supernatant precipitate were both resuspended in the same volume of 1x SDS Buffer (Laemmli Solution [Bio-Rad], 1 mM DTT) to create a concentration of 4 x 10^9^ CFU/ml, or an OD_600_ of 10 – typically 60-80 µl. Samples were boiled at 90 °C for 10 min to denature proteins before brief centrifugation.

For SDS-PAGE, 50 µl of each sample and Precision Plus Protein Standards (Bio-Rad) were then applied to a 4-20 % Protean TGX Stain Free Gel (Bio-Rad) before separation at 200V for 35 min. Gels were imaged on the Bio-Rad Gel Dock, then transferred to nitrocellulose membranes using the Mixed MW program on the TransBlot Turbo Transfer System. Membranes were blocked with EveryBlot buffer (Bio-Rad) before incubation overnight at 4 °C with an affinity-purified Rabbit polyclonal anti-ExoS primary antibody (from Dr. Arne Rietsch, 1:5000). After washing 3x with TBS-T (0.1 % Tween-20 in 1x TBS Buffer) for 5 min each, membranes were then incubated for 1 h at RT with Goat anti-Rabbit HRP-conjugated secondary antibody (ThermoFisher A16096, 1:1000). Membranes were again washed with TBS-T, then exposed to Clarity Western ECL substrate (Bio-Rad) for 5 min before chemiluminescence and colorimetric imaging.

### Statistical analysis

Data on bacterial depth from Imaris were exported into Microsoft Excel and GraphPad Prism 9 or 10 for visualization and analysis. The percent depth for each individual bacterium was considered a unique data point and were aggregated across all fields of view for each eye as one biological replicate. Data from 3-6 biological replicates for each condition were combined into a single column for graphing and shown as median ± interquartile range. For all *ex vivo* experiments, data did not pass normal distribution tests. The Kruskal-Wallis test with Dunn’s multiple comparisons or Kolmogorov-Smirnov test were used to compare groups. For growth curve comparisons, One-way ANOVA with Dunnett’s multiple comparisons or a paired Student’s t-Test were used. P < 0.05 was considered significant.

## Acknowledgements

Thanks to Dr. Alain Filloux (Imperial College London, UK) for originally providing *P. aeruginosa* wild-type strain PAO1F.

## Contributions

EJ, DS, NGK, AR, DE and SF designed the experiments; EJ, DS, NGK and AR performed the experiments; EJ, AR, AS, DS, NGK, DE and SF analyzed and interpreted the data; EJ, DE, AR and SF wrote the manuscript; DE and SF supervised the study.

## Supplemental Material

**Supplemental Figure S1. Comparison of corneal epithelium traversal by *P. aeruginosa* and its T3SS mutants.** Representative images of corneal epithelium traversal by wild-type PAO1F compared to the Δ*pscC*, Δ*popBD* and Δ*exoSTY* mutants color-coded for traversal depth.

**Supplemental Figure S2. EGTA induction of *P. aeruginosa* T3SS gene expression was similar between PAO1 and T3SS mutants.** OD_600_-normalized GFP signal from pJNE05 (P*exoS*) carried by PAO1F or its T3SS mutants in the translocon pore Δ*popBD*, known exotoxins Δ*exoSTY*, or the transcriptional repressor Δ*exsD* after 24 h growth in T3SS-induction media [TSB plus gentamicin (200 <u>μg/ml),</u> monosodium glutamate (100 mM), glycerol (1 %) and EGTA (2 mM)]. Similar P*exoS* induction, i.e. T3SS induction, was observed across wild-type and mutants. N = 3 separate growth curves per group. Mean +/- standard deviation, ns = not significant (One-way ANOVA with Dunnett’s Multiple Comparisons).

**Supplemental Figure S3. Rhamnose-induction of ExsA rescues *ex vivo* traversal.** (A) OD_600_- normalized GFP signal from pJNE05 (P*exoS*) carried by a Δ*exsA* mutant complemented with rhamnose-inducible *exsA* with or without inclusion of rhamnose (Rha) (2 %) in the growth medium [TSB with gentamicin (200 <u>μg/ml),</u> monosodium glutamate (100 mM) and glycerol (1%)] after 24 h of growth. N = 3 separate growth curves per group. Mean +/- standard deviation, ** P < 0.01 (Paired Student’s t-Test). (B) Traversal depth after *ex vivo* infection of Δ*exsA* with expression of *exsA* remaining off or induced by overnight growth with rhamnose (2 %). Data pooled from 3 eyes per strain. Error bars show the median with interquartile range. **** P < 0.0001 (Kolmogorov-Smirnov test).

