## Supplementary figures and images for "The *Pseudomonas aeruginosa* T3SS can contribute to traversal of an *in situ* epithelial multilayer independently of the T3SS needle"

### Supplemental Figure S1

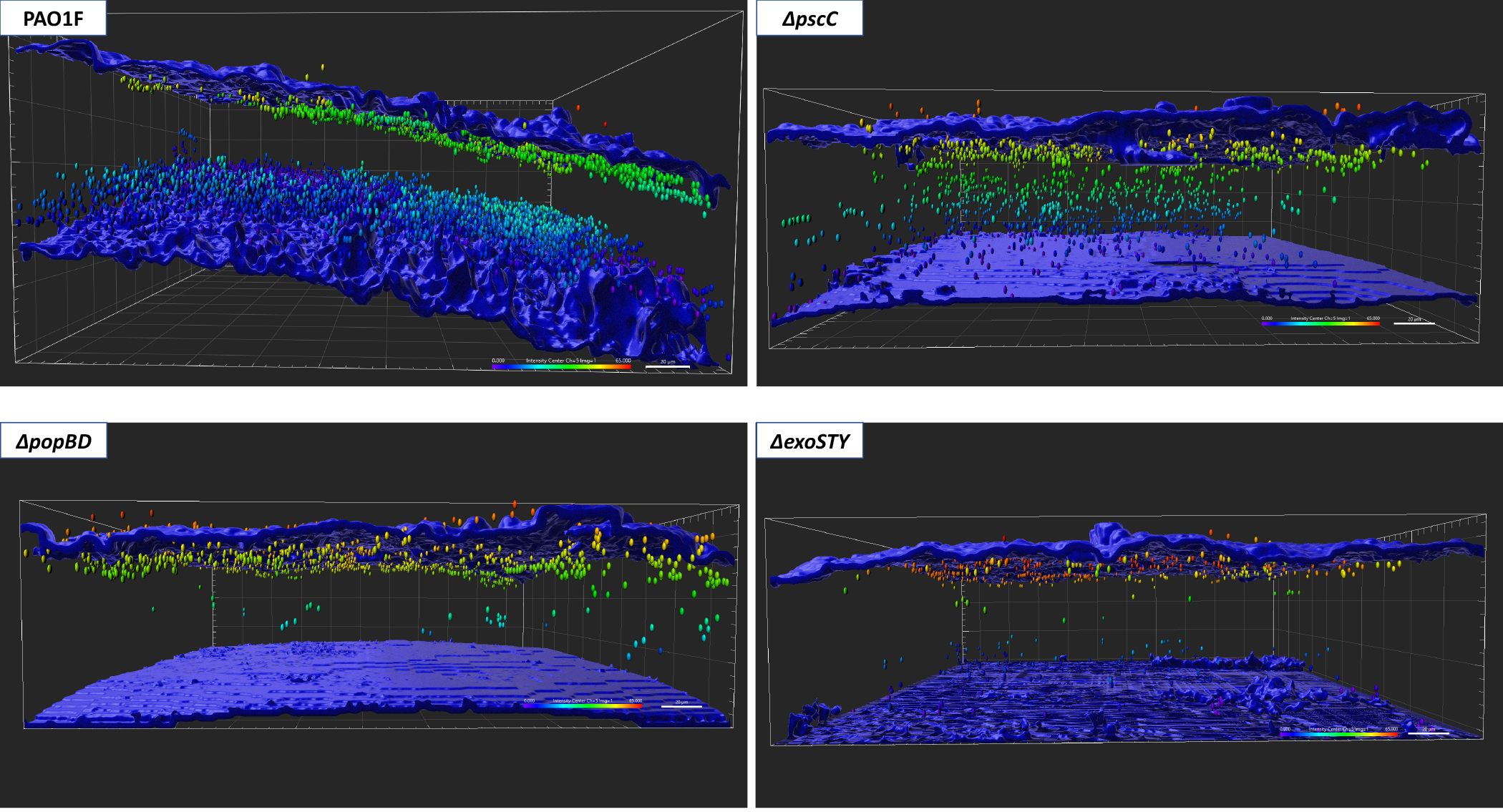

### Supplemental Figure S2

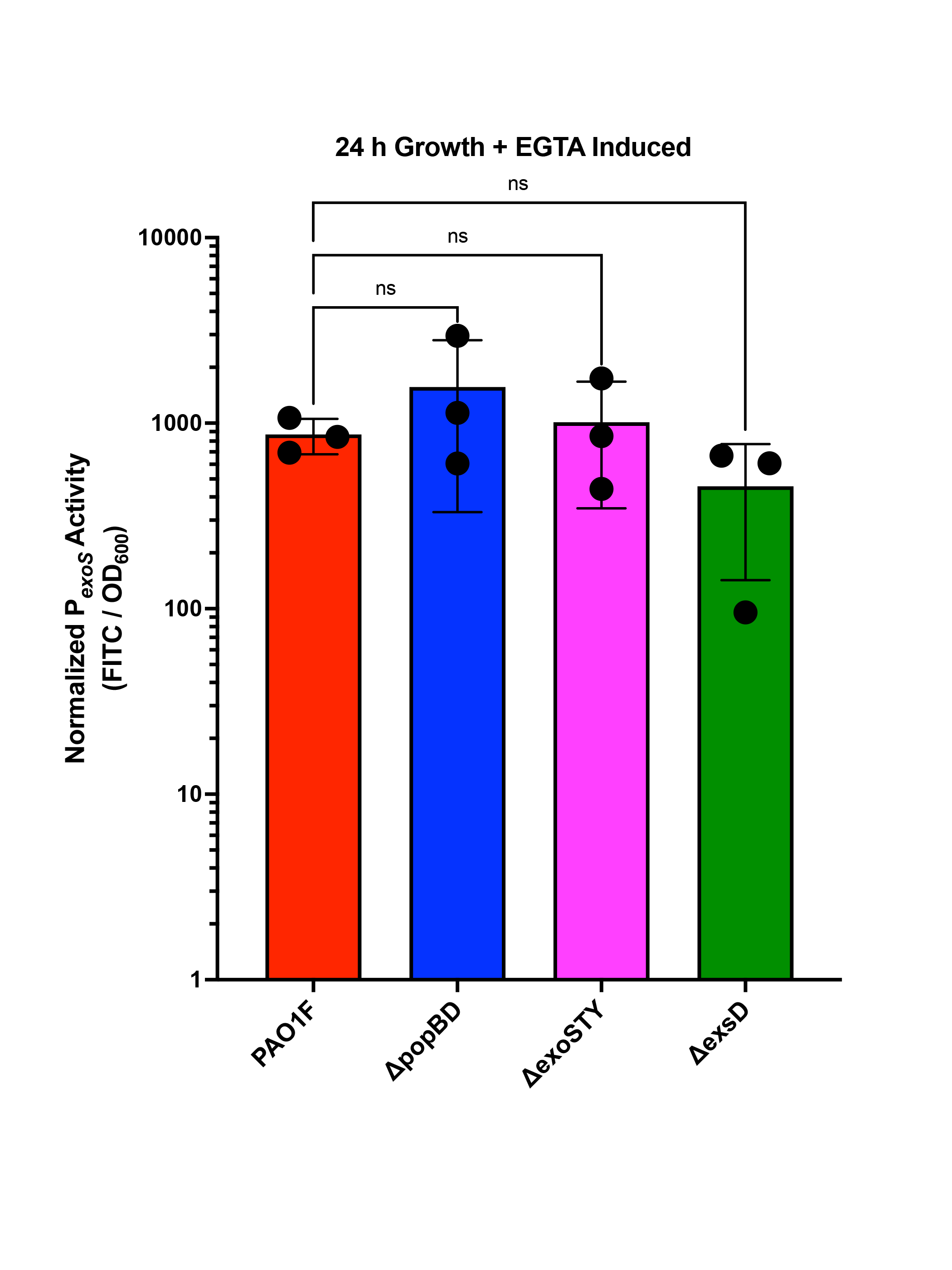

### Supplemental Figure S3

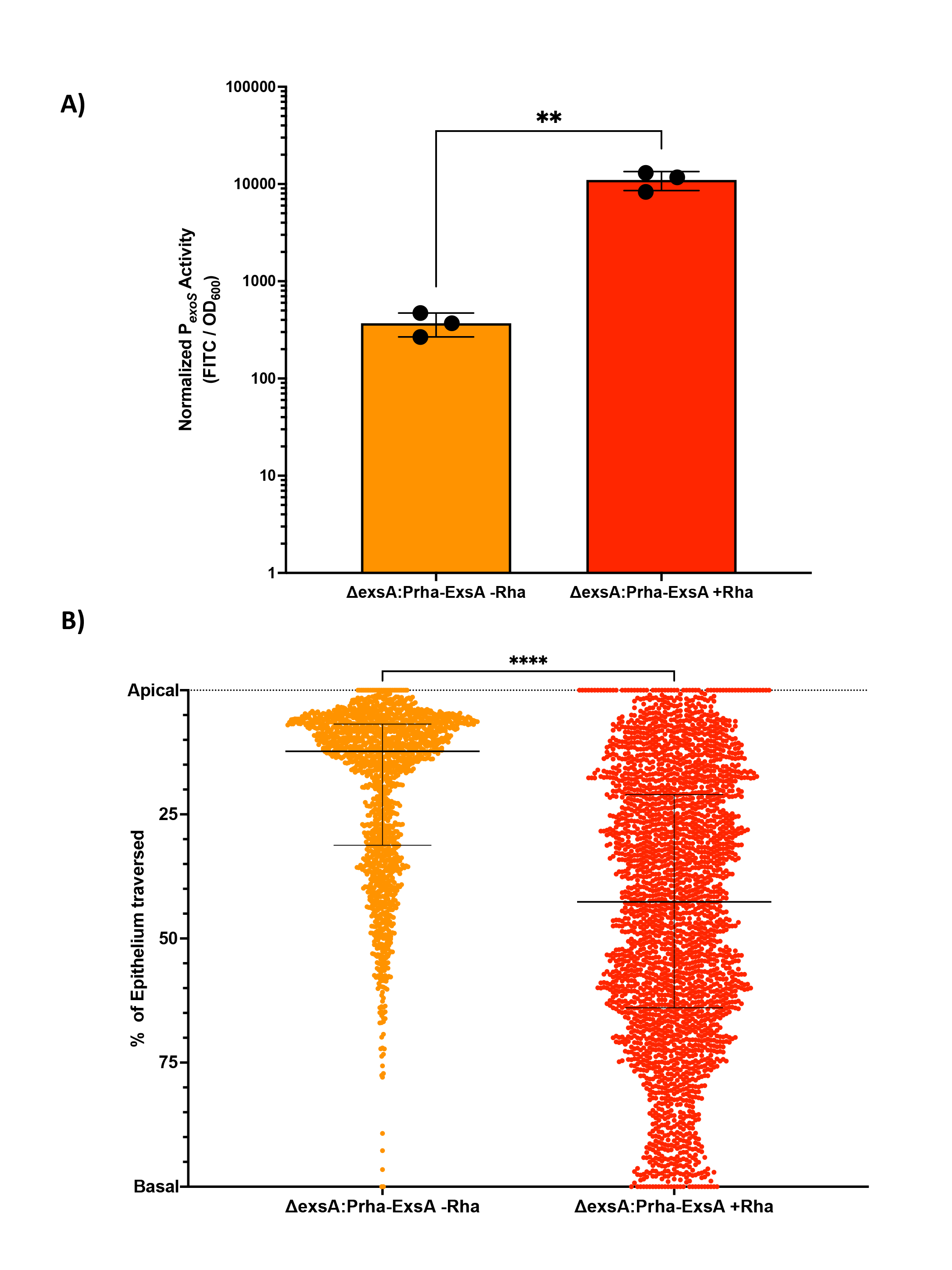
